# High Throughput Chromosome Conformation Capture identifies differential genome organization in virulent and avirulent strains of *Mycobacterium tuberculosis*

**DOI:** 10.1101/2022.11.10.515895

**Authors:** Mohit Mishra, Ajay Arya, Md. Zubbair Malik, Rakesh Bhatnagar, Seyed E. Hasnain, Shandar Ahmad, Rupesh Chaturvedi

**Affiliations:** School of Biotechnology, Jawaharlal Nehru University, New Delhi, 110067, India; School of Computational and Integrative Sciences, Jawaharlal Nehru University, New Delhi, 110067, India; Amity University, Jaipur, Rajasthan, India; Department of Biochemical Engineering and Biotechnology, Indian Institute of Technology Delhi, New Delhi, 110016, India; Special Center for System Medicine, Jawaharlal Nehru University, New Delhi, 110067, India; Nanofluidiks Pvt. Ltd. Jawaharlal Nehru University-Foundation for Innovation New Delhi, 110067, India; Department of Life Science, School of Basic Sciences and Research, Sharda University, Greater Noida, 201306, India; Department of Genetics and Bioinformatics, Dasman Diabetes Institute, Dasman, P.O. Box 1180, Kuwait city15462, Kuwait

**Keywords:** Hi-C, CID, CID boundaries, Gene expression

## Abstract

Recent studies have shown that three-dimensional architecture of bacterial chromatin plays an important role in gene expression regulation. However, genome topological organization in *Mycobacterium tuberculosis (M. tuberculosis)*, the etiologic agent of tuberculosis, remains unknown. On the other hand, exact mechanism of differential pathogenesis in the canonical strains of *M. tuberculosis* H37Rv and H37Ra remains poorly understood in terms of their raw sequences. In this context, a detailed contact map from a Hi-C experiment is a candidate for what bridges the gap. Here we present the first comprehensive report on genome-wide contact maps between regions of H37Rv and H37Ra genomes. We tracked differences between the genome architectures of H37Rv and H37Ra, which could possibly explain the virulence attenuation in H37Ra. We confirm the existence of a differential organization between the two strains most significantly a higher Chromosome Interacting Domain (CID) size in attenuated H37Ra strain. CID boundaries are also found enriched with highly expressed genes and with higher operon density in H37Rv. Furthermore, most of the differentially expressed PE/PPE genes were present near the CID boundaries in H37Rv and not in H37Ra. Collectively our study proposes a differential genomic topological pattern between H37Rv and H37Ra, which could explain the virulence attenuation in H37Ra.

## Introduction

Recent studies have unveiled that prokaryotic organisms feature very organized chromosome structures, which were hitherto discussed mainly for eukaryotic genomes (1–5). Among them, nucleoid-associated proteins (NAPs) help in maintaining the dynamic organization of the chromosome in the absence of histones in bacteria (6). Invention of chromosome conformation capture (3C) and its derivative techniques has opened up opportunities for investigating threedimensional structures of bacterial chromosomes and its long-range impact on gene expression regulation (7). Recent studies aimed at elucidating bacterial chromosome organization have provided enough evidence suggesting the presence of Chromosome Interacting Domains (CIDs) in bacteria, very similar to Topologically Associated Domains (TADs) found in eukaryotic organisms (1, 3, 5). Combination of 3C and Hi-C technology has made it possible to investigate comprehensive genome-wide interactions in some bacterial species such as *Escherichia coli (1), Caulobacter crescentus (3)*, and *Bacillus subtilis (4)*.

*Mycobacterium tuberculosis* is an extraordinary pathogen that latently infects almost one-third of the human population and also becoming refractile to treatment with rapidly developing multi-drug resistance (8, 9). H37Rv and its attenuated counterpart H37Ra both are derived from same parental strain H37 and have been widely used as laboratory strains for research aiming to understand *M. tuberculosis* pathogenesis (10). The H37Ra resembles the H37Rv genome in terms of gene order and gene content, however it is 8,445 bp longer because H37Ra has 21 deletions and 53 insertions compared to H37Rv strain (11). Major genetic changes are also caused by the differences in the repetitive sequences like IS6110 and the genes belonging to PE/PPE/PE-PGRS family (11). Despite various investigations over the years, the genetic causes for attenuation in H37Ra variant of *M. tuberculosis* remains poorly understood. The attenuated H37Ra strain is obviously expected to exhibit some alterations to either the genome or differential gene expression of virulence genes as compared to the virulent H37Rv. The *M. tuberculosis* genome is estimated to be more than 4 mega base long with over 4,000 protein-encoding genes, including 170 transcription factors (TFs), numerous DNA binding proteins, and many sigma factors each of which performs critical functions under various stress responses (12). How such a large number of functional entities are stuffed into this small genome remains an unresolved enigma.

By mapping a detailed contact organization in *M. tuberculosis*, using Hi-C, we aim to explore the genome organization in *M. tuberculosis*. Hi-C provides a two-dimensional map of complete threedimensional organization of chromatin in the form of pairwise genomic fragments data. In order to understand the three-dimensional genome organization in *M. tuberculosis*, in this study we have used exponentially growing mycobacterial culture. We performed Hi-C on both virulent and attenuated strain of *M. tuberculosis* and analyzed data by using Hi-C explorer tools (13). We constructed genome wide contact map at 10kb resolution and observed that the Ori and the midpoint are located at the two opposite poles of the chromosome structure. A differential CID organization was observed between virulent and attenuated strains with larger CIDs in attenuated strain H37Ra as compared to virulent counterpart H37Rv. We proposed that it could be a factor for virulence attenuation in H37Ra. We also observed that most of the CID boundaries were enriched with known highly expressed genes. Interestingly most of the genes belonging to PE-PPE family of genes with increased expression in H37Rv as compared to H37Ra are present near the CID boundaries in H37Rv. So collectively this differential CID organization in virulent and attenuated strain could indeed provide a novel way of transcriptional gene regulation and could be one of the mechanisms for attenuation.

## Results

### Comparative genome organization in virulent and attenuated *Mycobacterium tuberculosis* strains

To study chromosome organization in *M. tuberculosis*, we applied the Hi-C method to exponentially growing wild-type (WT) H37Rv and H37Ra cultures. A total of 75 million pairs of sequencing reads were generated for H37Rv and 110 million pairs of reads were generated for H37Ra (*SI Appendix*, Table S1). To analyze the contact information contained in them, we first mapped the resulting sequencing reads to the reference genome of H37Rv (NC 000962.3 NCBI) and H37Ra (NC_009525.1 NCBI) encompassing 4411532 bp and 4419977 bp, respectively. We obtained around 70 million high quality mapped reads in H37Rv and around 100 million in H37Ra. Further comprehensive sequence analysis identified around 19 and 40 million valid read pairs in H37Rv (26%) and H37Ra (41%), respectively for 3D genome construction. Intra long-range interactions (>= 20kb) revealed by Hi-C were much more frequent than Intra short-range interactions (< 20kb) in both H37Rv and H37Ra (*SI Appendix*, Table S1). The genome wide matrices were then constructed with valid Hi-C reads at 10 Kb resolution representing the final interaction frequencies (x, y), which reflects the relative contact frequency between bins x and y. We confirmed that biological replicates were highly correlated (Pearson correlation coefficient >0.92) in both H37Rv and H37Ra (*SI Appendix*, Fig. S1).

The generated H37Rv and H37Ra interaction matrix exhibits two prominent diagonals as consistent with other bacteria (Figure 1A). The two diagonals intersect each other at the center representing the terminus region (~2.2Mb) and at corners representing ori region (0 and 4.4Mb). This is consistent with the previous reports that the circular chromosome is organized in such a way that the origin and terminus occupies opposite poles of the cell and the chromosome is divided into left and right arm running along the axis (3, 14). The strong diagonal from top of left to bottom of right indicates that nearby loci were present on the same chromosomal arm and exhibit higher contact frequency. The less prominent diagonal bottom of left to top of right represents lower frequency contacts, i.e. those between opposite arms of the circular genome. Substantial distances at physical level separate these genomic loci pairs but the Hi-C data suggest that they were in close proximity, enabling their interactions in cellular context. We then compared the contact maps of H37Rv and H37Ra to examine the differences in their chromosome organization best studied at 10kb resolution in this work (Figure 1A). We observed a global similarity in the interaction map of H37Rv and H37Ra suggesting that the overall shape of chromosome remained conserved across these variants. Yet, critical differences were observed as discussed below.

**Figure 1:**
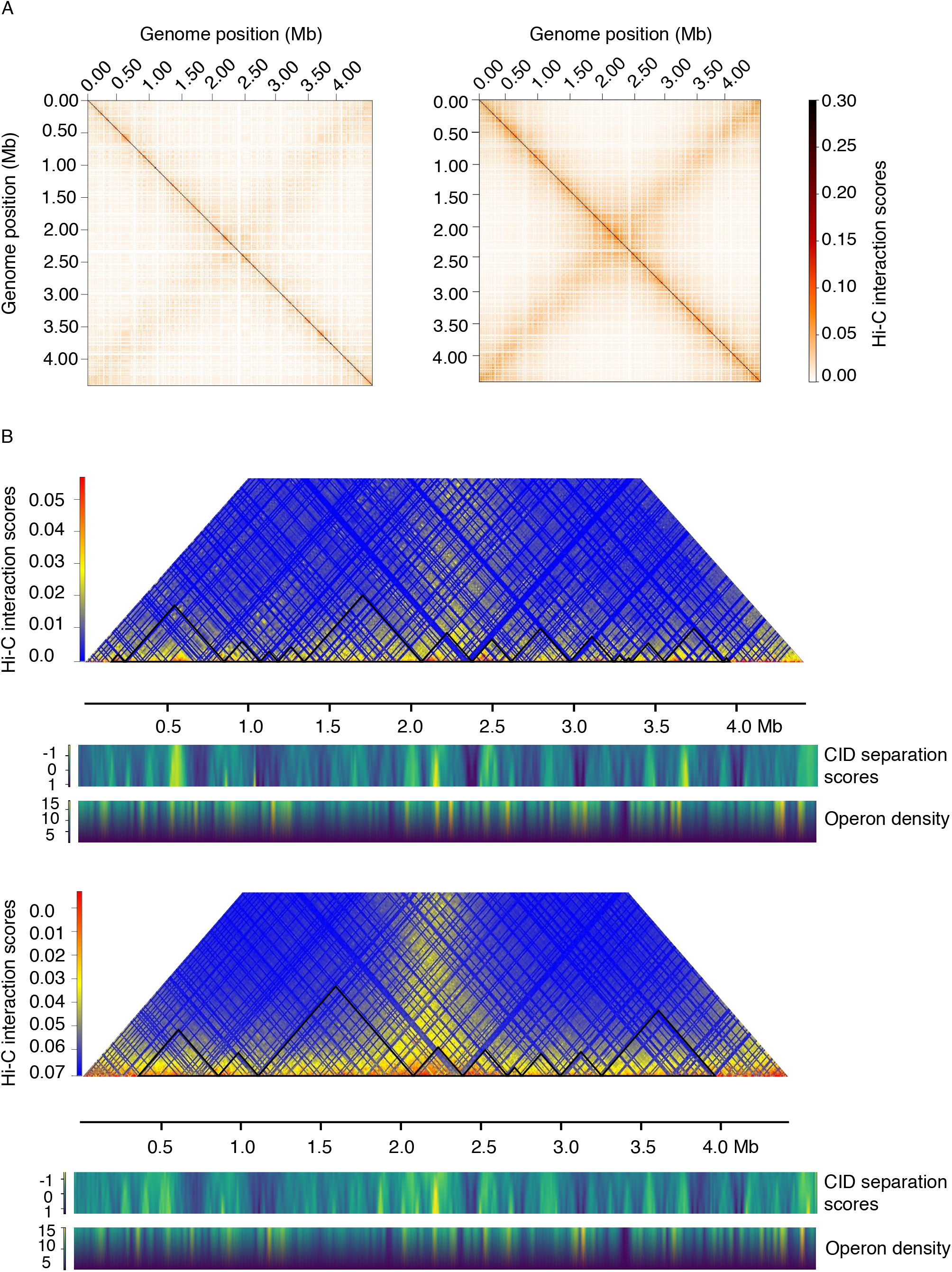
Comparative analysis of structural organization of circular chromosome in *Mycobacterium tuberculosis* strains H37Rv and H37Ra. A) Normalized BglII Hi-C contact map of *M. tuberculosis* strain H37Rv and H37Ra in exponential phase at a 10 kb resolution. The color of the contact map, from white to red, indicates the Log2 contact frequency. Axes indicate the genome position of each bin. B) Exemplary snapshot of identified CIDs in *M. tuberculosis* H37Rv and H37Ra. CID separation scores and operon densities are plotted below the CID Plot for both the strains.

### Virulent strain has higher number of Chromosome Interacting Domains than attenuated strain

In order to find the differences between the chromatin organization of CIDs in H37Rv and H37Ra, we employed domain caller software hicFindTAD (Methods) to detect the CIDs from corrected Hi-C interaction matrices generated at 10kb resolution. Despite a similarity between interaction maps of H37Rv and H37Ra at a global level, there was a considerable alteration in the number of CIDs as indicated by changes in their genomic positions (Figure 1B). Our analyses revealed that the genome of H37Rv is partitioned into a total of 15 CIDs (total genomic occupancy of 85.90%) with size ranging from 40kb to 720kb (*SI Appendix*, Table S2A), whereas the H37Ra chromosome comprised of 9 CIDs (total genomic occupancy of 81.70%) with size ranging from 90kb to 970kb (Figure 1B) (*SI Appendix*, Table S2B). Strikingly, we found that the median CID size was significantly higher in attenuated H37Ra (280 kb) as compared to virulent H37Rv (220 kb) (*SI Appendix*, Fig. S2A). While some of the CIDs were conserved in both H37Rv and H37Ra, others were altered as indicated by changes in their genomic positions. Besides the few unique CIDs (CID 1 in H37Ra and CID 1 and 2 in H37Rv) and similar CIDs (CID 2,4,5 and 8 of H37Ra with CID 3,7,8 and 10 of H37Rv), we observed that few larger CIDs in H37Ra (CID 3 and CID 9) were partitioned into two or more smaller CIDs in H37Rv (CID 4,5 and 6 and CID 11,12,13,14 and 15) respectively. Conversely, two CIDs (CID 6 and CID 7) in H37Ra coalesce into a larger ‘merged’ CID in H37Rv (CID 9). Collectively, these studies provide an indication of differential structural organization of CIDs in virulent and attenuated *M. tuberculosis* strains. To eliminate the possibility that the differential CID organization could be because of differences in total number of valid reads in H37Rv and H37Ra, we also generated contact maps and CID plots for H37Ra by using similar number of reads as H37Rv and confirmed the robustness of observed differences (*SI Appendix*, Fig. S3).

We further calculated the CID separation score for each CID boundary and plotted their summary for both H37Rv and H37Ra. We found that despite differential CID organization in virulent H37Rv and attenuated H37Ra strains, there was no significant difference in the strength of CID boundaries (*SI Appendix*, Fig. S2B) as indicated by their CID separation scores. We observed that operon density is higher at CID boundaries compared to within the CIDs (Figure 1B). Also CID separation scores and operon densities show a similar pattern in both H37Rv and H37Ra. However, newly created CID boundaries in region 3240kb-3960kb in H37Rv exhibit higher operon densities as compared to H37Ra. The *nrdHIEF2* operon, which includes *nrdH* (Rv3053c), *nrdI* (Rv3052c), *nrdE* (Rv3051c) and *nrdF2* (Rv3048c) plays an important role in chromosome duplication and DNA repair (15), and the alteration in expression of *nrdHIEF2* operon might impact growth and survival of *M. tuberculosis*. Interestingly *nrdHIEF2* operon is placed near the boundary in H37Rv within CID 13 (3350-3550) (59kb) whereas in case of H37Ra, this operon is placed within CID 9 around 177kb away from the boundary (*SI Appendix*, Fig. S4). Similarly The NADH dehydrogenase type I operon, encoded by nuoAN is present at 22kb from the CID boundary in H37Rv while in case of H37Ra this operon is present within the CID 9, 273kb away from the boundary.

### Chromatin loop formation exhibits differential preferences in right and left arm of the chromosome in H37Rv and H37Ra

Chromatin loops bring distant regulatory segments into close proximity, thereby affecting their transcription (16, 17). DNA loop formation in bacteria is attributed to the nucleoid-associated proteins (NAPs) such as H-NS, FIS and bacterial SMC proteins (6, 18, 19). To detect long-range contacts in both H37Rv and H37Ra, we employed hicDetectLoops program of Hi-C Explorer. HicDetectLoops can detect enriched interaction regions based on a strict candidate selection, negative binomial distributions and Wilcoxon rank-sum tests. The maximum genomic distance limits the candidate selection, which in–this study is 2MB. We identified 26 loops in H37Rv and 24 loops in H37Ra. Although we did not observe much difference between the number of loop formations in H37Rv and H37Ra, yet most of the long-range loops formed in H37Rv were present on the right arm of the chromosome i.e. from region 2 Mb to 4 Mb whereas in case of H37Ra most of the loops were formed in a region 0.2Mb to 2.5Mb (Figure 2A). We also observed that most of the loops were unique and very few loops were common in both the strains (Figure 2B) (Supplementary Table S3). The size of loops in H37Rv ranges from 10kb to 1790kb whereas it ranges from 30kb to 1900kb in H37Ra strain. However, the median loop size (840 kb) was higher in case of H37Ra as compared to H37Rv (515Kb) (Figure 2C).

**Figure 2:**
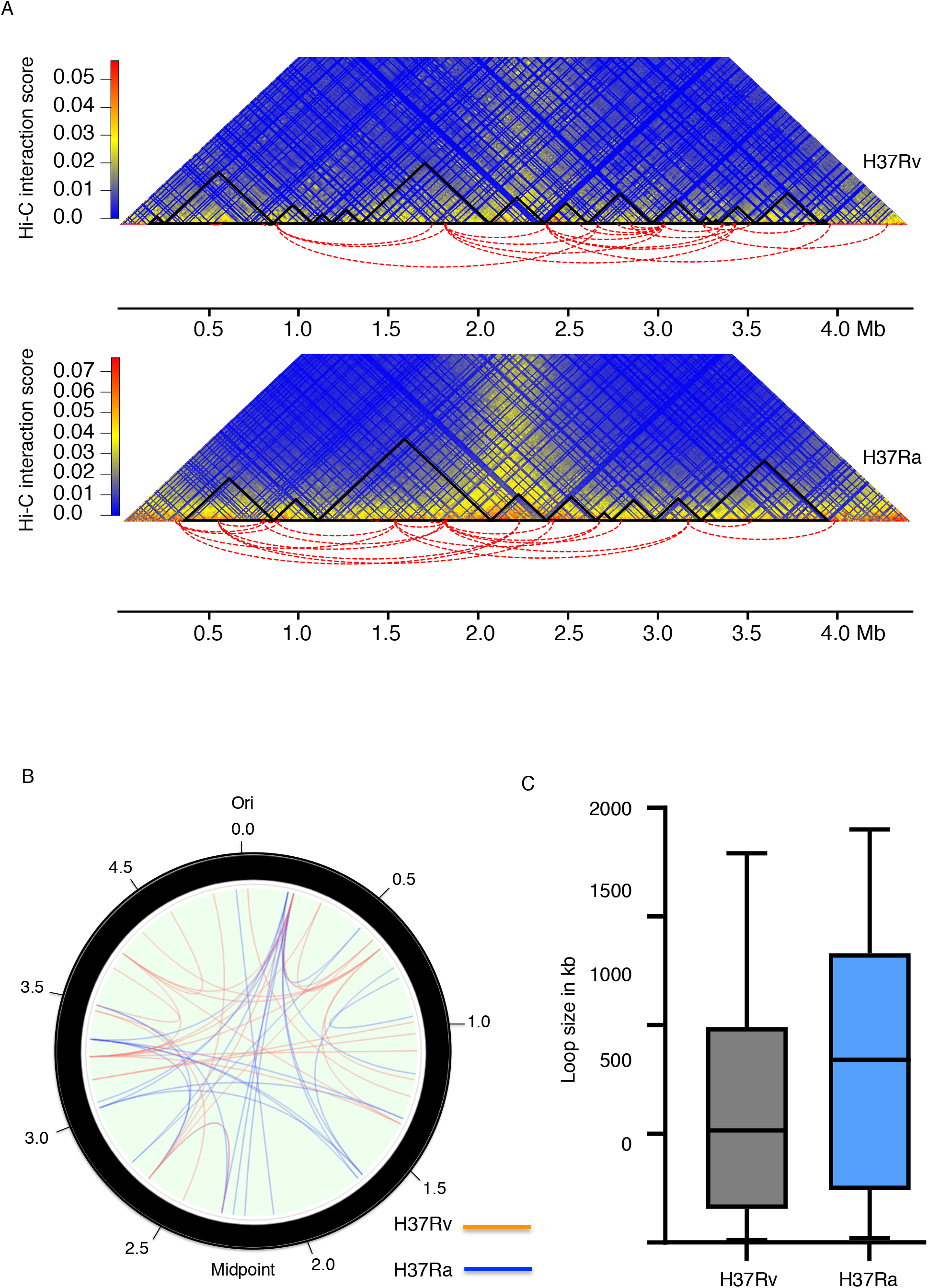
Chromatin loops identified in *M. tuberculosis* H37Rv and H37Ra genome. A) Long range Hi-C loops identified using hicDetectLoops program of Hi-C explorer in H37Rv and H37Ra. Arcs highlighted by red dotted lines represent the genomic positions of loops. B) Circos plot of intra-chromosomal interactions in H37Rv and H37Ra (Blue representing loops in H37Ra and orange in H37Rv). C) Comparison of loop size in H37Rv and H37Ra (Wilcoxon signed-rank test, P > 0.05).

### Creation of new CID boundaries in H37Rv in region 3250kb-3960kb corresponds to highly expressed genes in H37Rv as compared to H37Ra

To find out the gene expression profiles along the CIDs, we created a circular chromosomal map of *M. tuberculosis* H37Rv strain and marked the position of CIDs within it. We plotted average log FPKM score on the circular map of H37Rv genome (Figure 3A). We observed that most of the CID boundaries are enriched with highly expressed genes indicating a role of transcription in generation of CID boundaries in *M. tuberculosis*. Similarly, we plotted average log FPKM on circular map of H37Ra genome and observed that most of the CID boundaries in H37Ra were also enriched with highly expressed genes (*SI Appendix*, Fig. S5). This phenomenon seems to replicate highly expressed genes being located in nucleosome-free regions, widely observed in yeast and candida organisms (20). We further plotted GC content over the CID maps of both these strains and observed that even though both H37Rv and H37Ra contain a GC rich genome (65%), their CID boundaries were marked with low GC content (*SI Appendix*, Fig. S6). By comparative analysis of CID organization in H37Rv and H37Ra, we observed that CID 3 of H37Ra corresponds to CID 4,5 and 6 of H37Rv. Similarly, CID 9 of H37Ra corresponds to CID 11,12,13,14 and CID 15 of H37Rv (Figure 3B) leading to a functional segregation for transcriptionally active zones. This differential organization in region 3240kb-3960kb created new CID boundaries in H37Rv as compared to H37Ra. To understand whether the creation of new CID boundaries around the region 3240-3960kb was consistent with the presence of highly expressed genes, we plotted genes with higher gene expression in H37Rv as compared to H37Ra on CID map corresponding to region 3240-3960 kb for both the strains. (Figure 3C). We used microarray data of differential gene expression of H37Rv and H37Ra available at GEO database (ID GSE7539) (21). We found most of the genes like hupB, PPE50-PPE51, PE31, PPE-60 and LipF having higher gene expression in H37Rv were present near the newly created boundaries corresponding to region 3240kb - 3960 kb in H37Rv. HupB gene is overexpressed in H37Rv (fold change 1.2) as compared to H37Ra and this gene is located 105 kb away from the boundary of CID 9 in H37Ra but in case of H37Rv creation of a new boundary at

**Figure 3:**
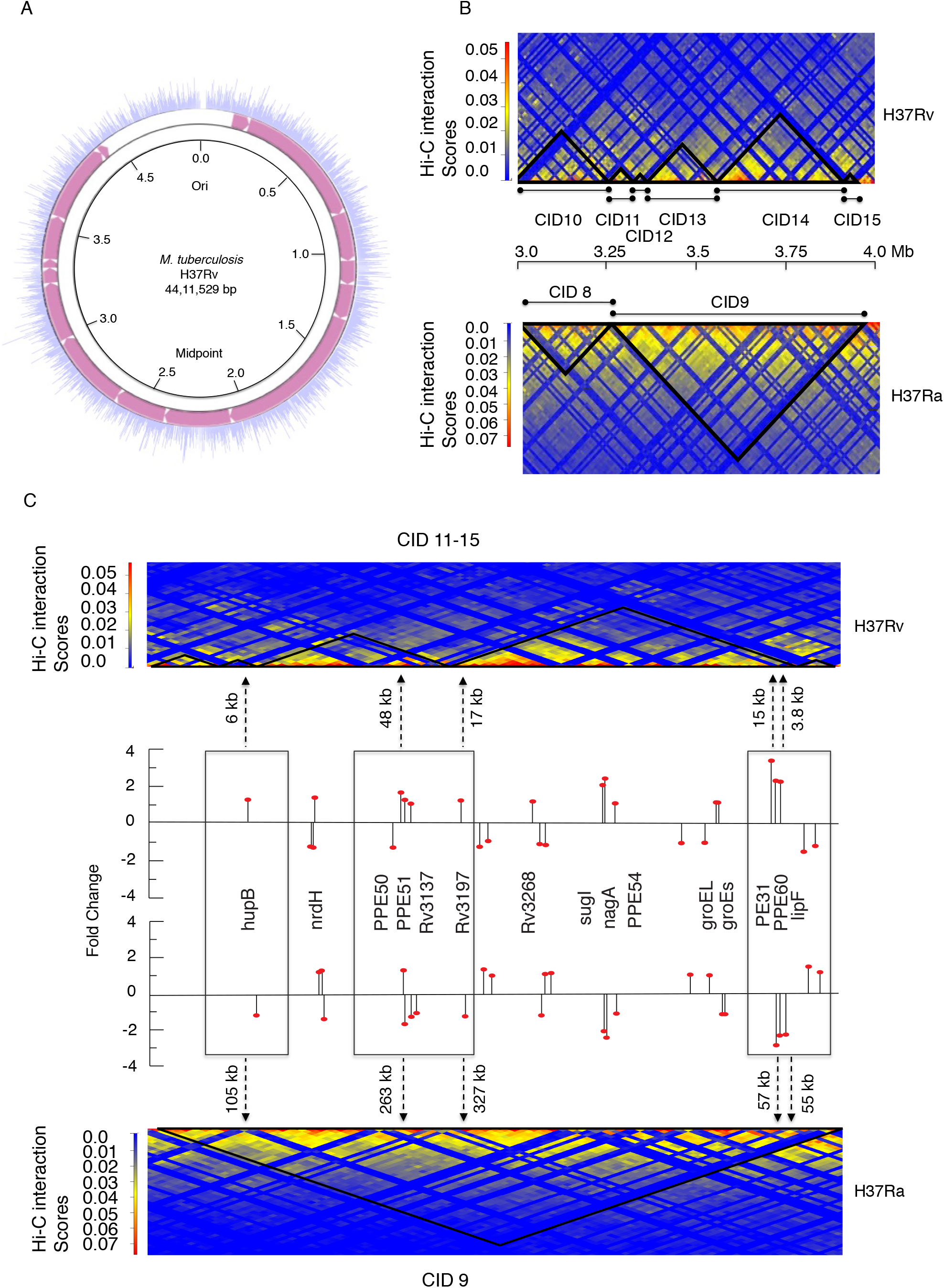
Highly expressed genes are present near the boundaries in H37Rv in a region corresponding to CID11-15. A) Circular chromosomal map showing the gene expression (FPKM values) across the chromosome of *M. tuberculosis* strain H37Rv with the positions of CIDs indicated by pink color. B) Comparison of presence of CIDs within the region 3240000-3960000 bp of *M. tuberculosis* chromosome. The upper and lower heat maps represents H37Rv and H37Ra, respectively. C) Plot showing genes with fold change expression (shown by black line) in H37Rv as compared to H37Ra on CID map corresponding to a region 3240000-3960000 bp for both the strains. The upper and lower heat maps represents H37Rv and H37Ra, respectively. Black dotted line with arrow indicates the genomic distances of genes from the CID boundaries.

CID 11(3240-3310kb) placed this gene 6 kb away from the boundary. Similarly, PPE50-PPE51 genes with increased expression in H37Rv (fold change 1.68 and 1.28, respectively) were placed at 48 kb near the CID 13 boundary as compared to 263kb away from the boundary in H37Ra. Similarly, PE31 and LipF genes that are overexpressed in H37Rv by a fold change of 3.3 and 2.5 respectively are placed at 15kb and 3.8kb from the CID boundary in H37Rv. We also plotted genes with differential gene expression in H37Rv as compared to H37Ra on CID map corresponding to region 1070-2070 kb for both the strains. It was observed that genes which are overexpressed in H37Rv such as pks3, fadD21, PE13 and PPE18 were present near the newly created boundary between CID 5 and 6 in H37Rv (*SI Appendix*, Fig. S7).

### CIDs corresponding to region 3240-3960kb in H37Rv shows enrichment for different pathways suggesting functional segregation of domains

To understand whether this differential CID organization is playing any role in terms of pathway enrichment, we carried out pathway enrichment analysis of CID 9 of H37Ra and corresponding CIDs 11-15 of H37Rv using ShinyGo tool (22). CID 9 of H37Ra did not show any enrichment however each of CID 11-15 of H37Rv showed enrichment for different biological pathways (*SI Appendix*, Fig. S8). For example CID 11 showed enrichment of pathways related to cell wall synthesis such as phthiocerol dimycocerosates (PDIMs), a group of complex lipids present in the *M. tuberculosis* cell envelope (23, 24). PDIM cluster is organized in a separate CID i.e. CID 11 in H37Rv. Similarly, CID 12 showed enrichment of genes related to carbohydrate metabolic process. Most of the pathways enriched in CID 13 are involved in oxidation–reduction processes including oxidative phosphorylation and ATP metabolic processes. CID 14 showed enrichment of genes related to nucleoside metabolic process as well as carbohydrate metabolism whereas CID 15 showed enrichment of cholesterol catabolism related genes. Collectively it was observed that CID 11-15 of H37Rv which corresponds to CID 9 of H37Ra showed enrichment of various different metabolic processes suggesting functional segregation of genes i.e. genes belonging to a particular metabolic process are clustered together.

### H37Rv CIDs in region 1070kb-2070kb and 3250kb-3960kb places differentially expressed PE/PPE genes near the boundaries

PE/PPE/PE-PGRS family of genes codes around 10% of the total genes of the mycobacterial genome (12, 25). The genes belonging to PE/PPE family are distributed throughout the mycobacterium genome and are implicated in most diverse functions such as virulence, host cell binding, and the immune system evasion (26). To explore the possibility of variability of PE/PPE genes in H37Rv and H37Ra, we plotted PE/PPE genes with differential gene expression in H37Rv and H37Ra on the CID map of both strains (Figure 4). We found that most genes belonging to PE/PPE family of proteins were preferentially present near the CID boundaries in H37Rv but less prominently so in H37Ra. Interestingly most of the PE/PPE genes present near the CID boundaries have higher gene expression in H37Rv as compared to those in H37Ra. PE13 and PPE18 gene pair is co-transcribed and are preceded by ESX gene pair. It has been shown that reduced expression of PE13 and PPE18 gene pair leads to attenuation of *M. tuberculosis* virulence (27). PE13-PPE-18 genes were present within interior region of CID 3 at 230kb away from the boundary. But in H37Rv this gene pair is moved to 1kb from the boundary region of CID 5. Similarly, PE31, a functionally important virulence gene, is overexpressed in H37Rv. This gene along with another PPE gene PPE60 is located 58kb away from the boundary of CID 9 in H37Ra but in H37Rv this gene is placed at 15kb from the boundary of CID 14. Another gene belonging to this family PPE18 appears to be present in a cluster with ESAT-6-like proteins (28) and is one of the highly expressed genes in *M. tuberculosis*. Interestingly, PPE18 harbored one of those deletions in H37Ra, and its deletion contributes to virulence attenuation of *M. tuberculosis in vivo* (11). Interestingly PPE-18, which is, located 240kb away from the boundary of CID 3 in H37Ra is moved near the boundary of CID 5 in H37Rv. So collectively these findings suggest that the creation of new CID boundaries in H37Rv resulted in positioning of these PE/PPE genes near the boundaries in H37Rv as compared to H37Ra.

**Figure 4:**
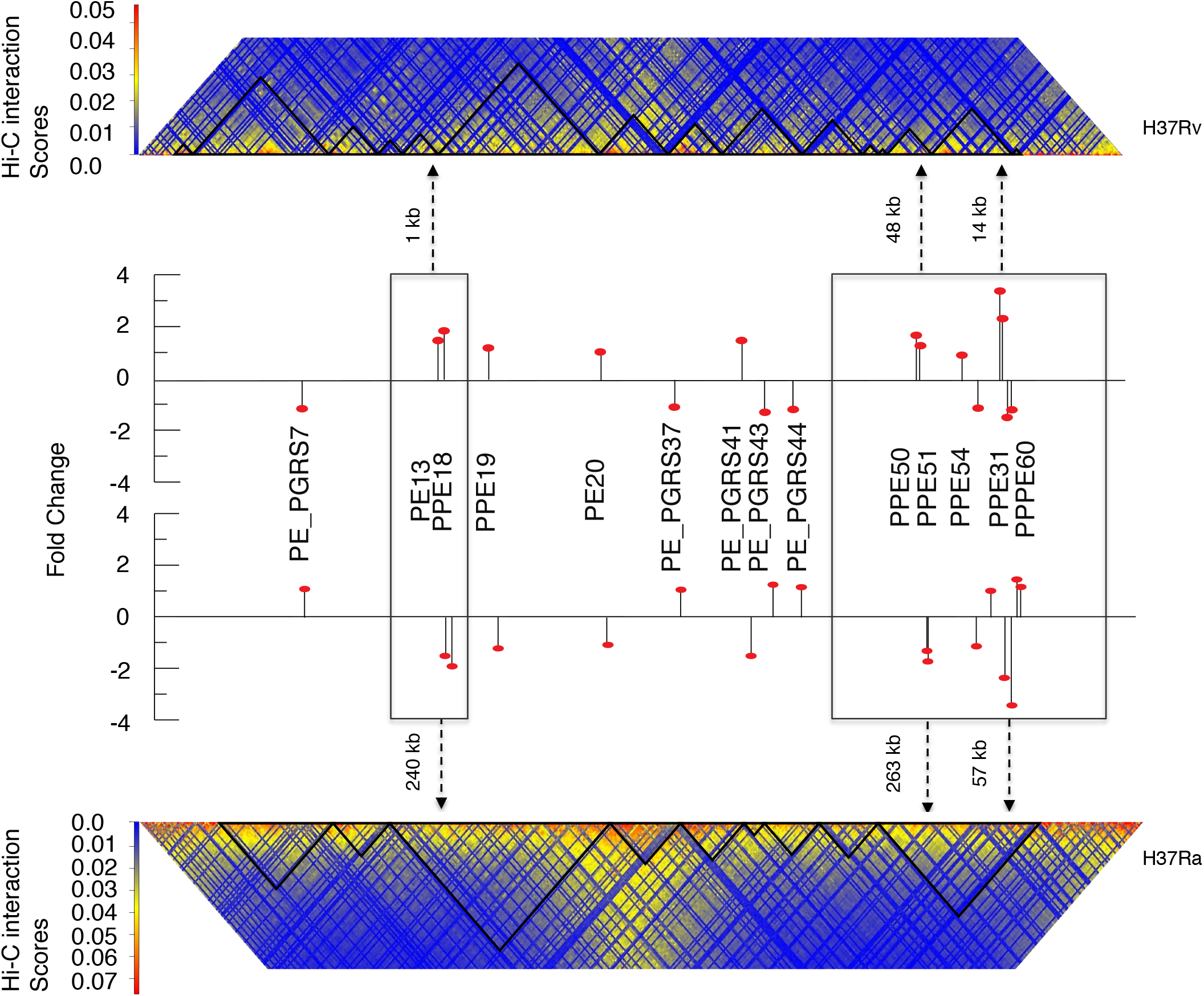
Differentially expressed PE and PPE genes are placed near CID boundaries in H37Rv as compared to H37Ra. Plot showing PE/PPE genes with fold change expression (shown by black line) in H37Rv as compared to H37Ra on CID map for both the strains. The upper and lower heat maps represents H37Rv and H37Ra, respectively. Black dotted line with arrow indicates the genomic distances of genes from the CID boundaries.

## Discussion

Chromosome conformation capture based methods such as Hi-C have been used in the past to study the bacterial genome organization and its role in transcriptional regulation. These studies in multiple bacteria such *E. coli, M. pneumonia, C. crescentus, and B. subtilis* have indicated that their genomes were organized in three-dimensional space as domains called chromatin interaction domains (CIDs) similar to topologically active domains (TADs) in eukaryotic cells (1–5). However, the comprehensive understanding of how the multilayer organization of chromatin regulates strain-specific transcriptional activity remains unclear. Furthermore, the Hi-C data have been collected for a relatively small number of organisms and none for a variant of *M. tuberculosis*. Current work presented on the two canonical strains of *M. tuberculosis* i.e. H37Rv and H37Ra, analyzed individually and comparatively provided much needed insights into how the genomic packing in the two variants occurs, specifically and non-specifically.

A key finding presented above is that H37Rv and H37Ra strains of *M. tuberculosis*, which are very much similar in order of gene content, exhibit critically different 3-dimensional organization of their genome. Several studies have previously shown that eukaryotic genomes were partitioned into TADs of size in the range of 200 kb to 1Mb (16, 17, 29–31). Our analysis allowed the detection of eukaryotic TAD-like domains (CIDs) for the first time in *M. tuberculosis*. CIDs in both *M. tuberculosis* strains are larger than those previously reported in other bacteria such as *C. crescentus, and B. subtilis* (3, 4). Our findings show that, a substantial number of CIDs were conserved between H37Rv and H37Ra suggesting a robust and meaningful three-dimensional topology in this family of bacteria across variants. Strikingly, however, we also found a number of CIDs that were appearing dynamic with structural alterations between H37Rv and H37Ra potentially providing for a previously unknown basis of their functional specificity. The major differences were observed in genomic regions corresponding to CID 3 (1100-2070 kb) and CID 9 (3250-3960 kb) of H37Ra. The corresponding genomic region in H37Rv showed the incidence of more numerous and smaller CIDs. The 710kb region in H37Rv appeared partitioned into five CIDs (CID 11-15) compared to the corresponding region in H37Ra. A study performed by Boya et al has earlier shown alteration in the structural organization of TADs during developmental transition from pre-pro-B to pro-B cell stage (32). They found that pro-B cells have a set of merged TADs generated due to coalesce of contiguous TADs present in pre-pro-B cells. However, the mechanism that regulates merging and splitting of CIDs remains to be understood. As both H37Rv and H37Ra have evolved from a same parental strain, this differential structural organization of CIDs during their evolution could be a possible factor for their differential pathogenesis. Presence of multiple insertions and deletion in H37Ra could be a possible driver of creation and abrogation of these CIDs during evolution.

Other than the numerosity of CIDs, nature and frequency of chromatin loops are the other topological characteristic of genomic organization. In eukaryotic genomes it has been shown that these loops bring distant regulatory segments into close proximity to regulate their transcription (16, 17). In eukaryotic cells CTCF protein and the protein complex cohesin have been implicated in the formation and maintenance of these chromatin loops, establishing CTCF as the master weaver of eukaryotic genome (33, 34). Long-range chromatin loop formation is poorly studied in bacterial genomes. Even though proteins like CTCF and cohesin are missing from them, they do contain several nucleoid-associated proteins (NAPs) such as H-NS, FIS and bacterial SMC proteins that have been shown to participate in bacterial DNA looping (6, 18, 19). In our study we did not observe significant differences between the number of long-range loops between H37Rv and H37Ra strains. However, interestingly we did observe that the loops were present on different arms of chromosome in virulent and avirulent strains of *M. tuberculosis*.

Previously CID boundary formation was correlated with the presence of highly expressed genes in *C. crescentus*. This way, the DNA is kept free of plectonemic loops during active transcription (3). In this study we have also observed that CID boundaries in both H37Rv and H37Ra were marked by the presence of highly expressed genes. Apart from highly expressed genes, we also observed that low GC content levels characterize CID boundaries. Similar reports in *E. coli* and *S. typhimurium* and more recently in *B. subtilis* suggested that domain loop boundaries were localized in AT-rich regions (35, 36). Apart from prokaryotic organisms, it has been observed in peanut genome too that TAD boundaries exhibit lower GC content as compared to TAD-interior region (37). Furthermore the active chromatin compartment in eukaryotic genome showed lower GC content and significantly higher transcription levels than inactive compartment (37). Our results also revealed that most of the differentially expressed genes between virulent H37Rv and avirulent H37Ra were present close to the boundaries particularly in H37Rv and not in H37Ra. This observation is most pronounced in the regions corresponding to 1070-2070kb and 3240-3960kb region in both H37Rv and H37Ra. We found that these results may explain some of the poorly understood experimental observations in published literature. For example HupB, an HU homologue was recently identified in mycobacteria and it is one of the most abundant nucleoid associated proteins. HupB gene is differentially expressed in H37Rv and H37Ra and present near the CID boundary in H37Rv. It has been previously reported that deletion of the hu1 and hu2 genes in C. crescentus, which encode the HU1 and HU2 proteins, significantly decreased short-range interactions but did not affect global chromosome organization (11, 18). The role of HupB protein in domain organization in mycobacterium strains still needs to be elucidated and the chromatin looping could be a potential reason for these observations.

Apart from placing highly expressed genes near the boundaries in H37Rv, the differences in CID organization within the region 3240-3960kb between H37Rv and H37Ra also account for different enrichment of pathways in CID 11-15 in H37Rv. For instance CID 11 in H37Rv showed enrichment of genes involved in cell wall synthesis (PDIM). Genes involved in PDIM biosynthesis are coded by a 70kb gene cluster present within the region 3240-3310 kb and interestingly the size of CID 11 (3240-3310) was also found to be 70kb (12). This was remarkable in the fact that all genes involved in a similar pathway are clustered within a CID. Also within this cluster, fadD26, ppsA–E, drrA–C and papA5 were also reported to form a single transcriptional unit (24). Similarly, CIDs 12,13,14 and 15 showed enrichments for different pathways which suggest that the formation of these smaller CIDs in H37Rv might be playing a role in segregating genes involved in different pathways.

PE/PPE/PE-PGRS family of genes contribute about one tenth of the coding capacity of *M. tuberculosis* and these genes are reported to be involved in various functions including virulence. Comparative genomic study between H37Rv and H37Ra discovered differences in genomic sequences of 35 PE/PPE/PE-PGRS genes. Furthermore, several PE/PPE/PE-PGRS genes found to be preferential “hot spots” for mutations (11). In our study, we found that most of PE/PPE genes with higher gene expression in H37Rv were placed near the boundaries in H37Rv.

Taking the above observations on CIDs and chromatin looping into account, we can conclude that even though H37Rv and H37Ra are very similar in DNA sequence, their three-dimensional structure differ significantly. One would wonder how the sequence and structure of bacterial genomes come together to perform the biological function, including virulence. In particular, it would be interesting to know how small changes in sequences may lead to large scale changes in CIDs and looping or local folding patterns of the DNA. This question becomes pertinent in view of topological changes between the two bacterial strains studied here in which appear to be driven by sequence level alterations. How much sequence-level changes are sufficient to introduce topological changes remains a question to be explored. On the other hand Hi-C provides a relatively lower resolution data and small and subtle changes in topologies that could occur at the local level in a sequence-dependent manner cannot be elucidates from this data. However, there is a body of evidence that shows that intrinsic sequence-dependent changes in double helical shape of the DNA can substantially impact transcriptional regulation, which can be explained only after translating the sequence signatures to their sequence-dependent DNA shapes (38). We have in the past shown that sequence-dependent conformational ensembles at the static and dynamic DNA shape level can fill the gaps in the knowledge of target specificity of transcription factors (38). Due to the pioneering works from other research groups in this direction, it is now well known that shape signatures contained in genomic sequences are critical factors to consider for understanding their specificities. Thus, long range DNA topological and local short range DNA shape changes seem to emerge as the fundamental pillars of understanding functional differences between genomes beyond their sequences, and the current work focuses on the former.

## Materials and Methods

### Growth conditions

*M. tuberculosis* strains were cultured in 7H9 medium supplemented with ADC enrichment (5% Albumin, 2% dextrose, 0.003% catalase and 0.85% sodium chloride) and 0.05% Tween 80. All the cultures were grown at 37°C with constant shaking at 200 rpm in a biosafety level 3 facilities. OD at 600 nm was measured to monitor growth.

### Hi-C Library preparation

All experiments were performed in BSL-3 facility. *H37Ra and H37Rv* chromatin was prepared as described previously with some modifications (12) (*SI Appendix*, Fig. S9). Briefly a total of 10^9^ cells were cross-linked with 1% formaldehyde and reaction was quenched with glycine. Crosslinked cells were lysed using lysozyme and lysed cells were then digested using BglII enzyme. After successful digestion, Biotin Fill-in was performed and cells were ligated under dilute conditions. DNA was isolated using phenol chloroform method after proteinase K treatment. After removing biotin from unligated ends, Hi-C library was prepared using NEBNext^®^ Ultra™ II DNA Library Prep Kit for Illumina and final DNA library was sequenced in the HiSeq Illumina platform. Sequencing reads were aligned, mapped and filtered before generating a genome-wide contact matrix at 10kb resolution using Hi-C explorer pipeline.

### Generation of Hi-C contact map

HiCExplorer is a set of programs to process, normalize, analyze and visualize Hi-C and cHi-C data, available on GitHub. Before using HiCExplorer to build a Hi-C contact matrix paired-end reads were mapped, aligned and filtered. Only valid Hi-C reads were used to generate contact maps at 10kb resolution. Then raw contact maps were normalized and corrected using KR correction method, which balances a matrix using a fast balancing algorithm introduced by (39).

### Identification of chromosome interacting domains

We used hicFindTADs program of HiCExplorer, which uses a measure called CID-separation score to identify the degree of separation between the left and right regions at each Hi-C matrix bin to detect the CIDs. We kept minDepth at 30000bp, and max depth at 60000bp with step size equal to 10000bp.

### Loop detection

A program of Hi-C Explorer called hicDetectLoops was used for loop detection. hicDetectLoops detect enriched interaction regions (peaks / loops) based on a strict candidate selection, negative binomial distributions and Wilcoxon rank-sum tests. Experimental procedures including Hi-C library preparation, sequencing and data analysis are described in details in Method S1.

## Acknowledgments

The authors would like to thank the Jawaharlal Nehru University facility and all funding agencies for supporting us.

## References

1. Cagliero C, Grand RS, Jones MB, Jin DJ, & O’Sullivan JMJNar (2013) Genome conformation capture reveals that the Escherichia coli chromosome is organized by replication and transcription. 41(12):6058–6071.

2. Lioy VS, et al. (2018) Multiscale structuring of the E. coli chromosome by nucleoid-associated and condensin proteins. 172(4):771–783. e718.

3. Le TB, Imakaev MV, Mirny LA, & Laub MTJS (2013) High-resolution mapping of the spatial organization of a bacterial chromosome. 342(6159):731–734.

4. Wang X, Brandão HB, Le TB, Laub MT, & Rudner DZJS (2017) Bacillus subtilis SMC complexes juxtapose chromosome arms as they travel from origin to terminus. 355(6324):524–527.

5. Trussart M, et al. (2017) Defined chromosome structure in the genome-reduced bacterium Mycoplasma pneumoniae. 8(1):1–13.

6. Wang W, Li G-W, Chen C, Xie XS, & Zhuang XJS (2011) Chromosome organization by a nucleoid-associated protein in live bacteria. 333(6048):1445–1449.

7. Sati S & Cavalli GJC (2017) Chromosome conformation capture technologies and their impact in understanding genome function. 126(1):33–44.

8. Chakaya J, et al. (2021) Global Tuberculosis Report 2020–Reflections on the Global TB burden, treatment and prevention efforts. 113:S7–S12.

9. Chakaya JM, et al. (2020) Programmatic versus personalised approaches to managing the global epidemic of multidrug-resistant tuberculosis. 8(4):334–335.

10. Steenken Jr W & Gardner LJArot (1946) History of H37 strain of tubercle bacillus. 54(1):62–66.

11. Zheng H, et al. (2008) Genetic basis of virulence attenuation revealed by comparative genomic analysis of Mycobacterium tuberculosis strain H37Ra versus H37Rv. 3(6):e2375.

12. Cole S, et al. (1998) Deciphering the biology of Mycobacterium tuberculosis from the complete genome sequence. 396(6707):190–190.

13. Wolff J, et al. (2018) Galaxy HiCExplorer: a web server for reproducible Hi-C data analysis, quality control and visualization. 46(W1):W11–W16.

14. Umbarger MA, et al. (2011) The three-dimensional architecture of a bacterial genome and its alteration by genetic perturbation. 44(2):252–264.

15. Dawes SS, et al. (2003) Ribonucleotide reduction in Mycobacterium tuberculosis: function and expression of genes encoding class Ib and class II ribonucleotide reductases. 71(11):6124–6131.

16. Rao SS, et al. (2014) A 3D map of the human genome at kilobase resolution reveals principles of chromatin looping. 159(7):1665–1680.

17. Liu C & Weigel DJGb (2015) Chromatin in 3D: progress and prospects for plants. 16(1):1–6.

18. Dillon SC & Dorman CJJNRM (2010) Bacterial nucleoid-associated proteins, nucleoid structure and gene expression. 8(3):185–195.

19. Dame RT, Tark-Dame M, & Schiessel HJMm (2011) A physical approach to segregation and folding of the Caulobacter crescentus genome. 82(6):1311–1315.

20. Ozonov EA & van Nimwegen EJPcb (2013) Nucleosome free regions in yeast promoters result from competitive binding of transcription factors that interact with chromatin modifiers. 9(8):e1003181.

21. Lee JS, et al. (2008) Mutation in the transcriptional regulator PhoP contributes to avirulence of Mycobacterium tuberculosis H37Ra strain. 3(2):97–103.

22. Ge SX, Jung D, & Yao RJB (2020) ShinyGO: a graphical gene-set enrichment tool for animals and plants. 36(8):2628–2629.

23. Rens C, Chao JD, Sexton DL, Tocheva EI, & Av-Gay YJM (2021) Roles for phthiocerol dimycocerosate lipids in Mycobacterium tuberculosis pathogenesis. 167(3):001042.

24. Camacho L, Constant P, Raynaud C, Lanéelle M, & Triccas JJJBC (A., Gicquel, B., Daffé, M., and Guilhot, C.(2001) Analysis of the phthiocerol dimycocerosate locus of Mycobacterium tuberculosis. 276:19845–19854.

25. Fishbein S, Van Wyk N, Warren R, & Sampson SJMm (2015) Phylogeny to function: PE/PPE protein evolution and impact on M ycobacterium tuberculosis pathogenicity. 96(5):901–916.

26. Akhter Y, Ehebauer MT, Mukhopadhyay S, & Hasnain SEJB (2012) The PE/PPE multigene family codes for virulence factors and is a possible source of mycobacterial antigenic variation: perhaps more? 94(1):110–116.

27. Goldstone RM, Goonesekera SD, Bloom BR, Sampson SLJI, & immunity (2009) The transcriptional regulator Rv0485 modulates the expression of a pe and ppe gene pair and is required for Mycobacterium tuberculosis virulence. 77(10):4654–4667.

28. Gey van Pittius NC, et al. (2006) Evolution and expansion of the Mycobacterium tuberculosis PE and PPE multigene families and their association with the duplication of the ESAT-6 (esx) gene cluster regions. 6(1):1–31.

29. Gibcus JH & Dekker JJMc (2013) The hierarchy of the 3D genome. 49(5):773–782.

30. Lieberman-Aiden E, et al. (2009) Comprehensive mapping of long-range interactions reveals folding principles of the human genome. 326(5950):289–293.

31. Dixon JR, et al. (2012) Topological domains in mammalian genomes identified by analysis of chromatin interactions. 485(7398):376–380.

32. Boya R, et al. (2017) Developmentally regulated higher-order chromatin interactions orchestrate B cell fate commitment.

33. Li Y, et al. (2013) Characterization of constitutive CTCF/cohesin loci: a possible role in establishing topological domains in mammalian genomes. 14(1):1–12.

34. Zuin J, et al. (2014) Cohesin and CTCF differentially affect chromatin architecture and gene expression in human cells. 111(3):996–1001.

35. Noom MC, Navarre WW, Oshima T, Wuite GJ, & Dame RTJCB (2007) H-NS promotes looped domain formation in the bacterial chromosome. 17(21):R913–R914.

36. Marbouty M, et al. (2015) Condensin-and replication-mediated bacterial chromosome folding and origin condensation revealed by Hi-C and super-resolution imaging. 59(4):588–602.

37. Zhang X, et al. (2021) Chromatin spatial organization of wild type and mutant peanuts reveals high-resolution genomic architecture and interaction alterations. 22(1):1–21.

38. Andrabi M, et al. (2017) Predicting conformational ensembles and genome-wide transcription factor binding sites from DNA sequences. 7(1):1–16.

39. Knight PA & Ruiz DJIJoNA (2013) A fast algorithm for matrix balancing. 33(3):1029–1047.

