## Supplement File for "High Throughput Chromosome Conformation Capture identifies differential genome organization in virulent and avirulent strains of *Mycobacterium tuberculosis*"

#### Figure S1

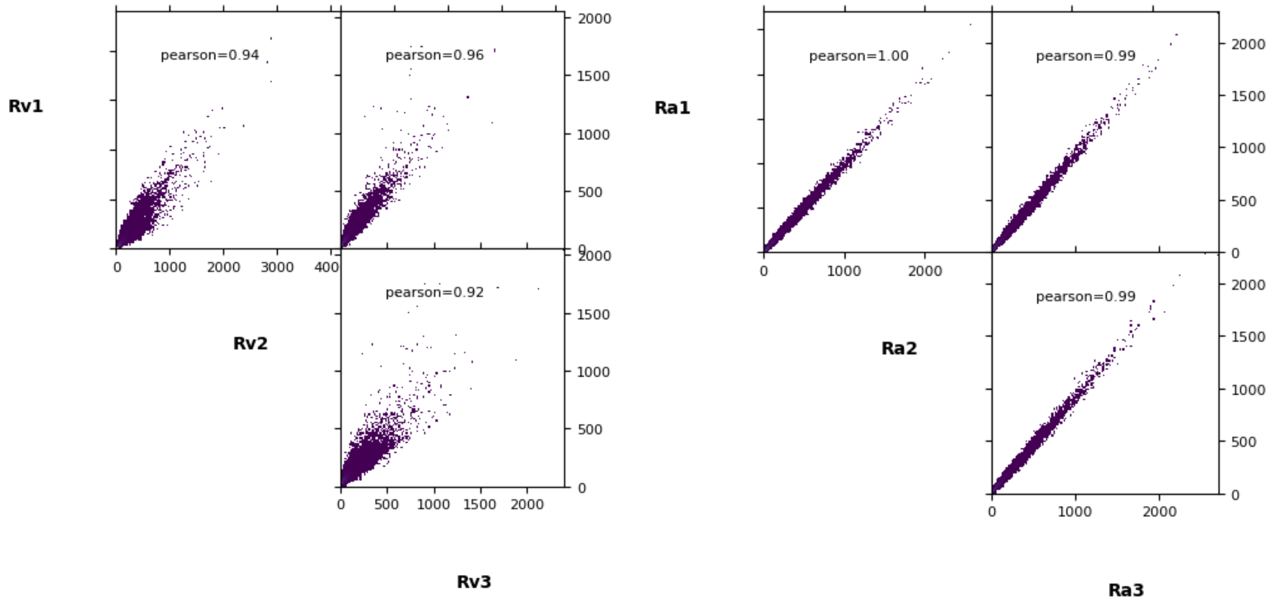


#### Figure S2

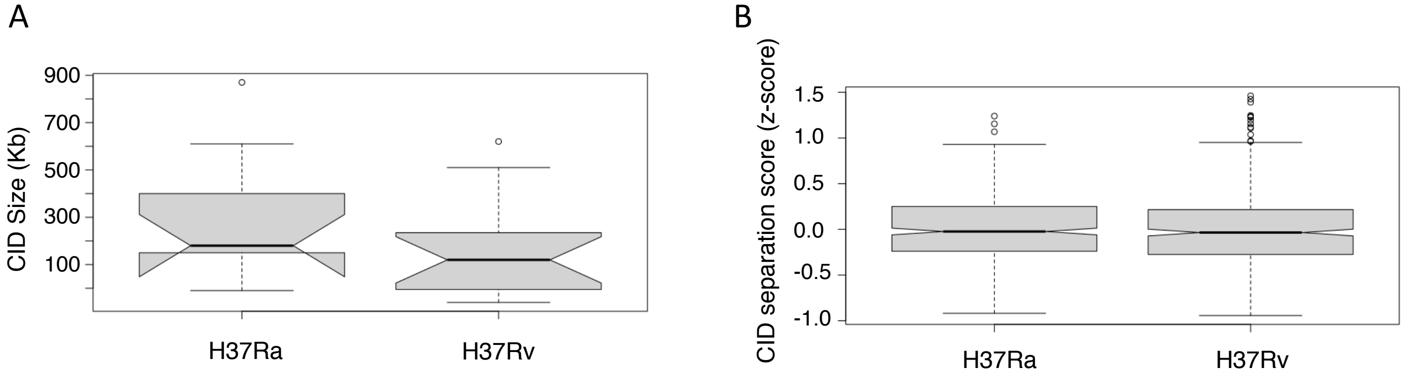

**Figure S3**

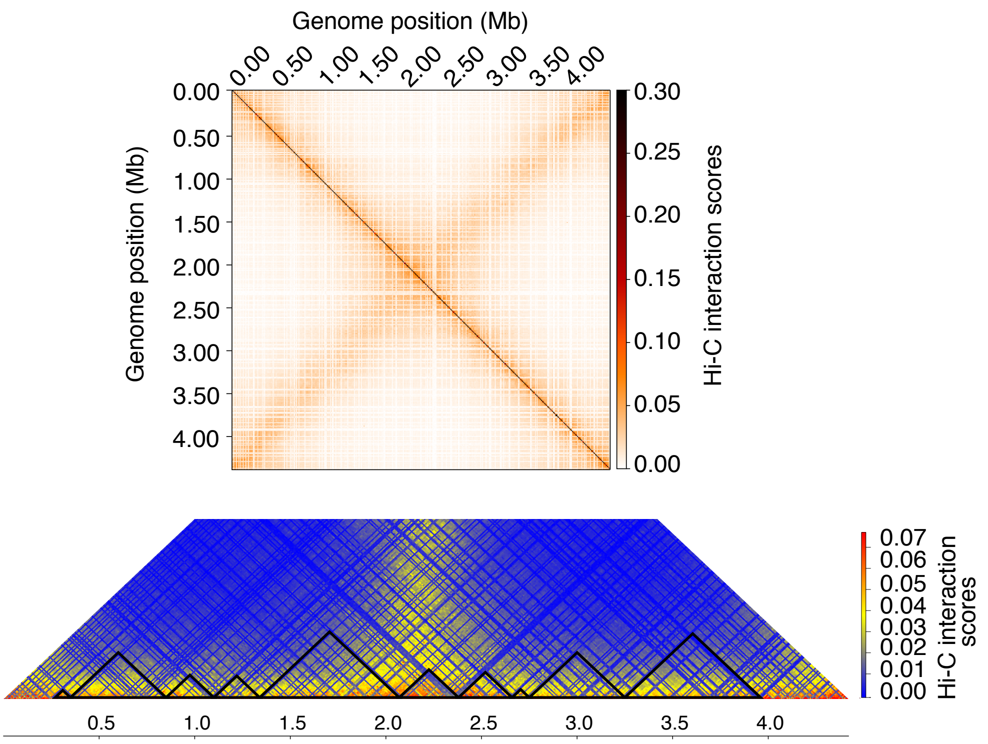


**Figure S4**

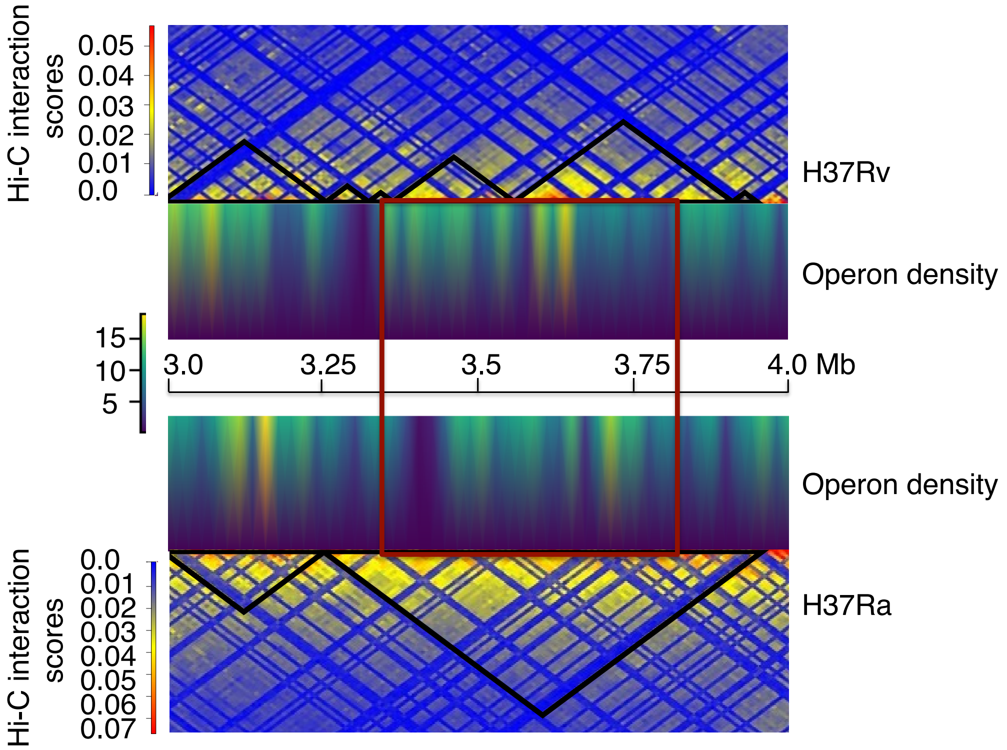

**Figure 5S**

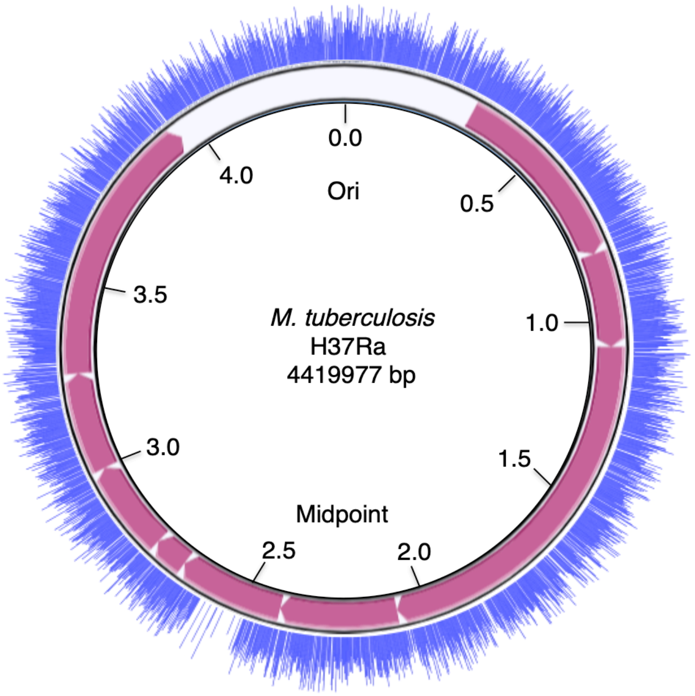


**Figure 6S**

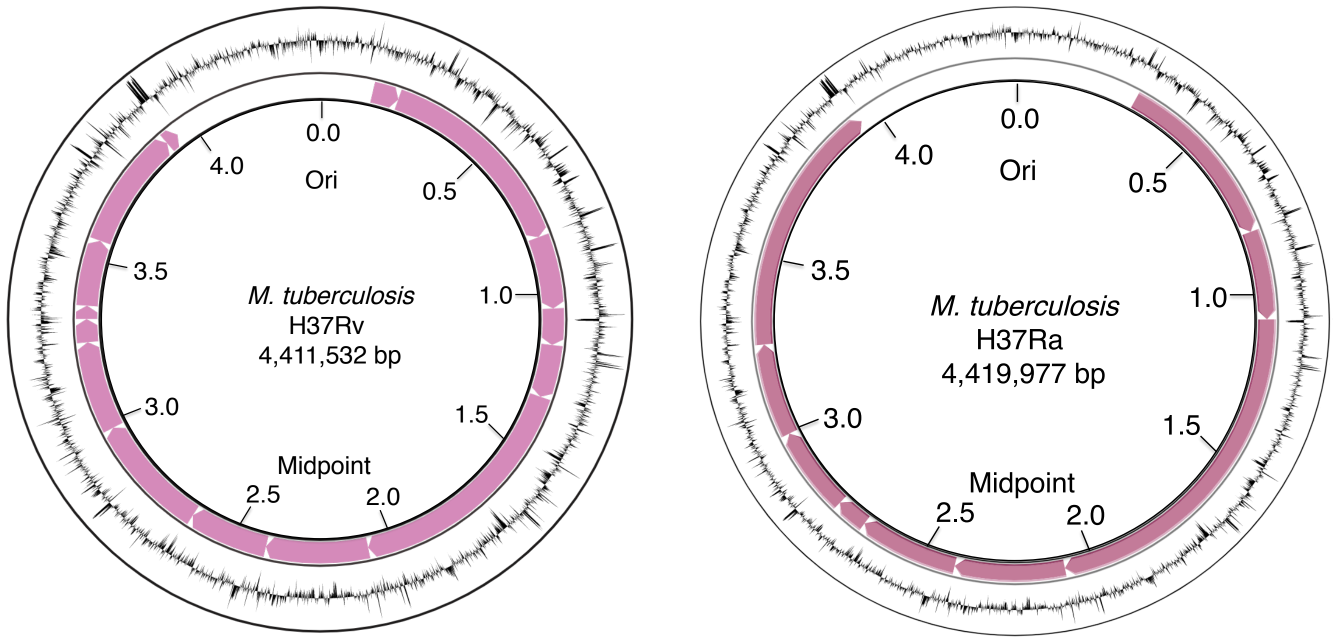

**Figure 7S**

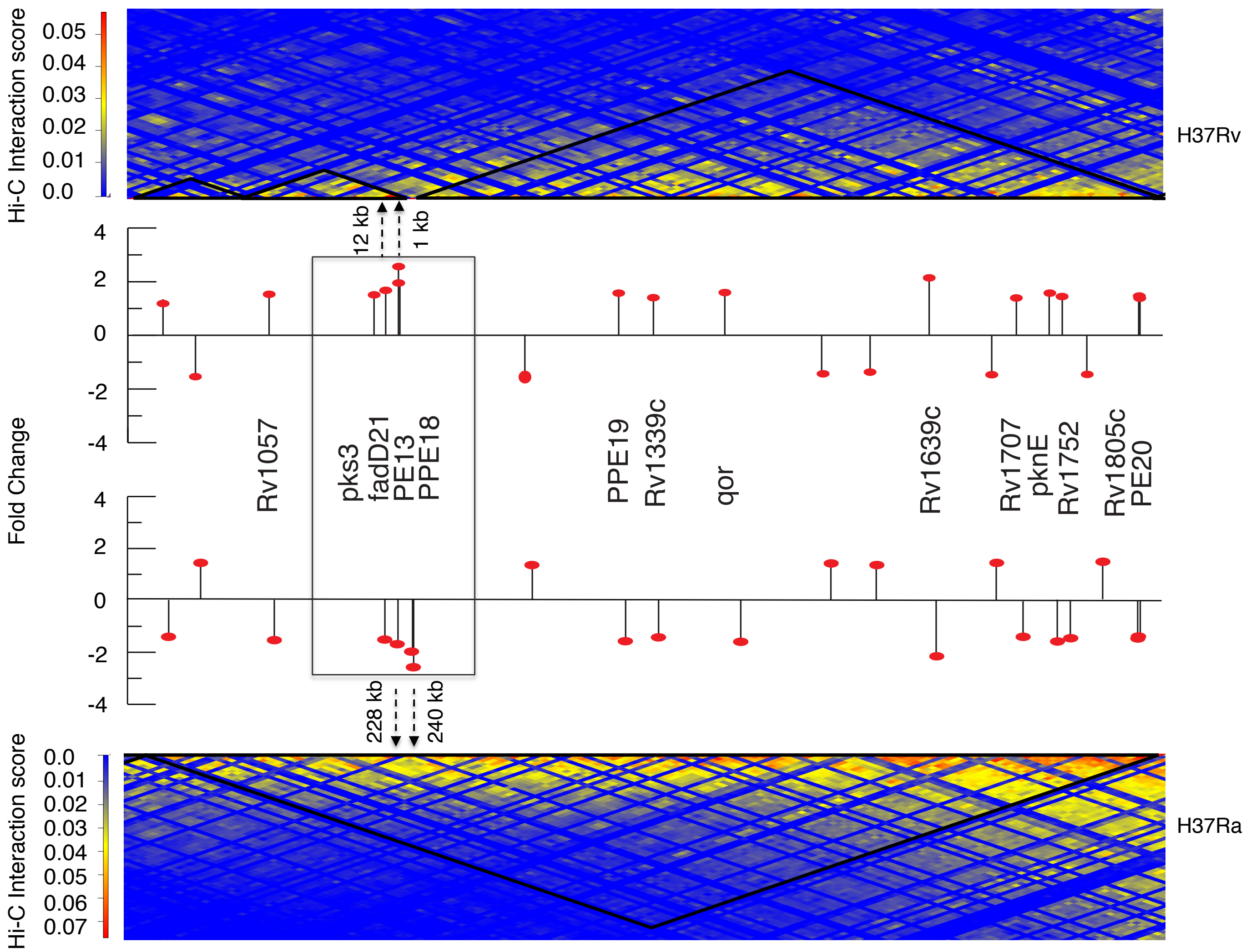
Hi-C Interaction score
0.05
0.04
0.03
0.02
0.01
0.0
H37Rv
12 kb
1 kb
Fold Change
4
2
0
-2
-4
Rv1057
pks3
fadD21
PE13
PPE18
PPE19
Rv1339c
qor
Rv1639c
Rv1707
pknE
Rv1752
Rv1805c
PE20
228 kb
240 kb
Hi-C Interaction score
0.0
0.01
0.02
0.03
0.04
0.05
0.06
0.07
H37Ra

### Figure S8

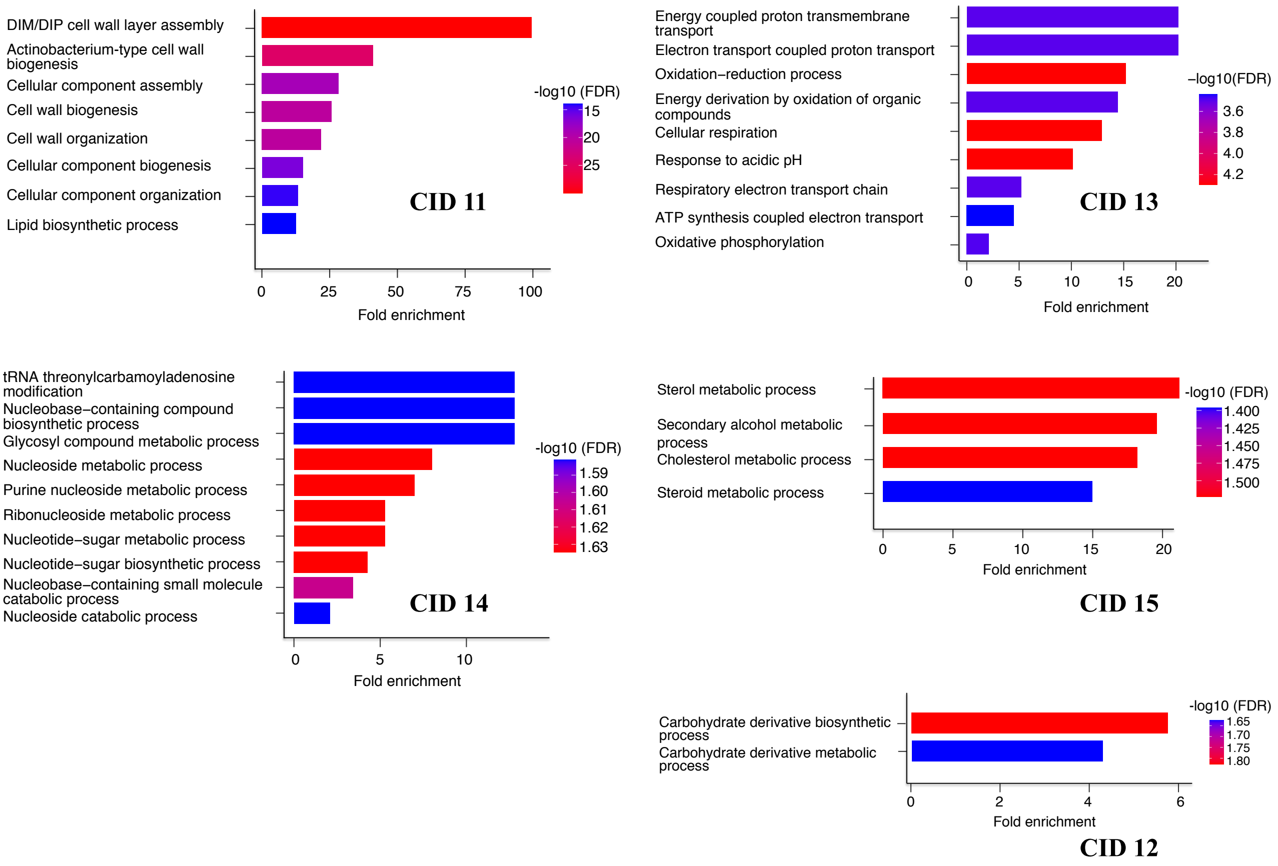

DIM/DIP cell wall layer assembly
Actinobacterium-type cell wall biogenesis
Cellular component assembly
Cell wall biogenesis
Cell wall organization
Cellular component biogenesis
Cellular component organization
Lipid biosynthetic process
0
25
50
75
100
Fold enrichment
-log10 (FDR)
15
20
25
CID 11
Energy coupled proton transmembrane transport
Electron transport coupled proton transport
Oxidation−reduction process
Energy derivation by oxidation of organic compounds
Cellular respiration
Response to acidic pH
Respiratory electron transport chain
ATP synthesis coupled electron transport
Oxidative phosphorylation
0
5
10
15
20
Fold enrichment
−log10(FDR)
3.6
3.8
4.0
4.2
CID 13
tRNA threonylcarbamoyladenosine modification
Nucleobase−containing compound biosynthetic process
Glycosyl compound metabolic process
Nucleoside metabolic process
Purine nucleoside metabolic process
Ribonucleoside metabolic process
Nucleotide−sugar metabolic process
Nucleotide−sugar biosynthetic process
Nucleobase−containing small molecule catabolic process
Nucleoside catabolic process
0
5
10
Fold enrichment
-log10 (FDR)
1.59
1.60
1.61
1.62
1.63
CID 14
Sterol metabolic process
Secondary alcohol metabolic process
Cholesterol metabolic process
Steroid metabolic process
0
5
10
15
20
Fold enrichment
-log10 (FDR)
1.400
1.425
1.450
1.475
1.500
CID 15
Carbohydrate derivative biosynthetic process
Carbohydrate derivative metabolic process
0
2
4
6
Fold enrichment
-log10 (FDR)
1.65
1.70
1.75
1.80
CID 12

**Figure S9**

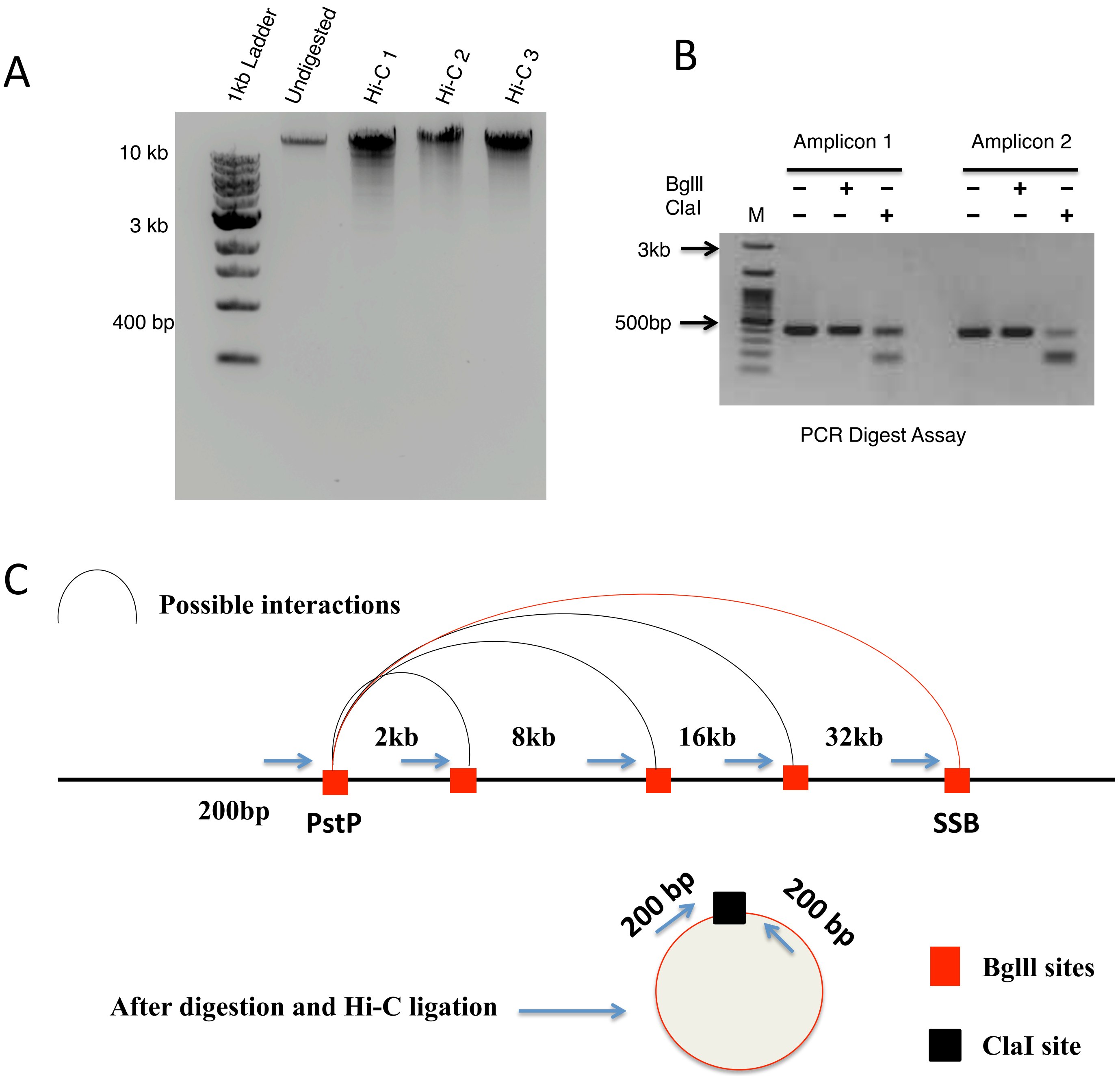
A
1kb Ladder
Undigested
Hi-C 1
Hi-C 2
Hi-C 3
10 kb
3 kb
400 bp
B
Amplicon 1
Amplicon 2
BglII
ClaI
M
– + –
– – +
– + –
– – +
3kb
500bp
PCR Digest Assay
C
Possible interactions
2kb
8kb
16kb
32kb
200bp
PstP
SSB
200 bp
200 bp
After digestion and Hi-C ligation
BglII sites
ClaI site

**Supplementary Figures:**

**Fig. S1. Pearson correlation between Hi-C replicates**

Pairwise scatter plots comparing interactions between three biological replicates in both H37Rv (Rv1, Rv2 and Rv3) and H37Ra (Ra1, Ra2, Ra3).

**Fig. S2. Comparison of CID size and CID separation scores in virulent and attenuated strains of *M. tuberculosis***

(A) Box plot showing the median CID size (kb) in attenuated H37Ra and virulent H37Rv strains.
(B) Box plot showing the CID separation score in attenuated H37Ra and virulent H37Rv strains.

**Fig. S3. Contact map and CID map generated in H37Ra using similar number of reads as H37Rv**

Normalized BglII Hi-C contact map and snapshot of identified CIDs in *M. tuberculosis* strain H37Ra. The color of the contact map, from white to red, indicates the Log2 contact frequency. Axes indicate the genome position of each bin.

**Fig. S4. Comparison of Operon density between H37Rv and H37Ra strains**

CID plots showing Operon density distribution in H37Rv and H37Ra in a region from 3 to 4 Mb. Red box highlights the region showing region involving CID 12,13 and 14 of H37Rv.

**Fig. S5. Gene expression profile of *M. tuberculosis* H37Ra strain**

Plot of log FPKM scores (blue lines) on Circular chromosome map of H37Ra with marked CIDs (represented by pink arrows).

**Fig. S6. Plot of GC content on CID maps of H37Rv and H37Ra**

Circular maps of H37Ra and H37Rv with marked CIDs (represented by pink arrows). GC scores are marked on the CID on an outer ring represented by black lines.

**Fig. S7. Plot showing differentially expressed genes with fold change on CID maps of H37Rv and H37Ra**

Plot showing genes with fold change expression (shown by black line) in H37Rv as compared to H37Ra on CID map corresponding to a region from 1070000 to 2070000 bp for both the strains. The upper and lower heat maps represents H37Rv and H37Ra, respectively. Black dotted line with arrow indicates the genomic distances of genes from the CID boundaries.

**Fig.S8. Go analysis of CIDs 11-15 in H37Rv**

Bar plot showing enrichment of Go terms in CID 11, 12, 13, 14 and 15 in *Mycobacterium tuberculosis* strain H37Rv.

**Fig. S9. Quality control of Hi-C libraries**

(A) Integrity of Hi-C libraries was monitored by resolving Hi-C templates (100 ng) on a 0.8% agarose gel. All templates were visualized as a tight band of a size larger than 10 kb.

(B) End fill-in efficiency of Hi-C libraries was monitored by amplifying a ligation junction formed by two BgIII restriction sites separated by a distance of 16 kb (amplicon 1) and 32 kb (amplicon 2). In Hi-C library successful fill-in and ligation of Bglll sites creates ClaI site, which is used to assess end fill-in efficiency of Hi-C libraries.

(C) A schematic representation of designing of unidirectional primers for PCR digest assay.
